# Diversity of mosquitoes (Diptera: Culicidae) collected in different types of larvitraps in an Amazon rural settlement

**DOI:** 10.1101/2020.06.23.166694

**Authors:** Jessica Feijó Almeida, Heliana Christy Matos Belchior, Claudia María Ríos-Velásquez, Felipe Arley Costa Pessoa

## Abstract

Anthropogenic environments provide favorable conditions for some species, which are especially suitable for mosquitoes that present eclecticism at the moment of choice for the site of oviposition. The present study, the diversity of mosquitoes in plastic container, bamboo internode, and tire placed in forest, the forest edge, and peridomicile environments in a rural settlement area was assessed. Eighteen sampling points were chosen, delimited by a buffer 200 m, placed in environments: forest, forest edge, and peridomicile. In each environment, larvitraps were installed, separated by a minimum distance of 7 m and 1 m from the ground. A total of 10,131 immature mosquitoes of 20 species were collected. The most abundant species was *Culex urichii* (29.5%), followed by *Trichoprosopom digitatum* (27.1%), and *Cx*. (Melanoconion) sp. (10,4%). There was a difference in the composition of immature mosquito populations between larvitraps (p<0.0005), and the plastic container hosted a greater diversity of species, whereas tires presented a greater abundance of individuals. The forest, forest edge, and peridomicile environments were also significantly different with regard to diversity of immature mosquito populations (p<0.0010). The forest edge was the environment with the greatest diversity of species, followed by the peridomicile and forest environments. In the forest and peridomicile, plastic container larvitraps had the greatest diversity, whereas the forest edge tire presented the largest number of individuals. Further, tire larvitraps collected the largest number of individuals in all environments. We identified 10 species associated with the bamboo internode and tire. We also observed the preference of species for artificial larvitraps, such as the plastic container and tire, even in wild environments. These artificial objects may represent a risk factor for the population living in this region, as all vector species found in the study were present in plastic containers and tires.

## Introduction

Insects are the most diverse of all animal classes on the planet, and Brazil is the country with the greatest insect diversity, with estimates of around 400-500 thousand known species and most of these inhabiting the Amazon forest [1–3]. Anthropic activity has affected some insect species, especially dipterian vectors such as sand flies, biting midges, and mosquito populations [4–6]. Mosquitoes are vulnerable to changes in environment and climate caused by deforestation and land use. Environmental changes affect the distribution of Culicidae, leading to the increased abundance of some species and a decreased abundance of others. Inevitably, the dynamics of disease transmission by mosquitoes are also affected [7,8].

Rural settlements are established in places that are favorable for human-vector interaction due to their proximity to extensive forest areas and the lack of adequate infrastructure for the residents who live there. In the Amazon region, precarious assistance by the government in rural settlements and the habits of local residents have led to an increase in cases of malaria and arboviruses [9,10].

A study carried out in a settlement in Amazonas, Brazil found a high seroprevalence of the Mayaro virus in residents, including people who did not enter the forest, such as children and women, suggesting that this arbovirus is also transmitted by species other than the main vector, *Haemagogus janthinomys* Dyar [11]. It was also observed that, in the settlement, adult *Ochlerotatus serratus* (Theobald), *Psorophora cingulata* (Fabricius), *Hg. tropicalis* Cerqueira and Antunes mosquitoes had been naturally infected with the *Oropouche* virus, in addition to the typical acrodendrophilous species captured in the soil. Further, a greater diversity of species was reported in the forest edge environment when compared to the forest and peridomicile [12].

Mosquitoes are characterized by their flexible use of a wide variety of breeding sites, from larges found at the soil level, to small pools of water stored in leaves in the canopy of trees. Man-made objects are also perceived as potential breeding sites for mosquitoes [13]. Immature *Limatus durhamii* Theobald mosquitoes, a species naturally infected with the *Guama* virus, can develop in breeding sites ranging from tree holes to landfill percolation tanks [14,15]. Immature forms of *Anopheles* species have been recorded in artificial containers [16,17], including *An. darlingi* Root, which was found in an artificial lagoon in the urban region of Manaus in the state of Amazonas [18].

Studies of immature mosquitoes captured in artificial breeding sites located in Amazonian environments are scarce. However, some authors [13,19–23] have indicated that artificial breeding sites are preferred by immature mosquitoes of various species.

Due to the importance of sanitary conditions and the ecological evolution that has led to the use of water containers for breeding by female mosquitoes, this work aimed to identify mosquito species that colonize different types of artificial breeding places, located in environments with different levels of anthropization within the state of Amazonas.

## Materials and Methods

### Study site and sample design

The study was carried out in the Rio Pardo rural settlement, Presidente Figueiredo Municipality, State of Amazonas, Brazil (Fig 1). The settlement is surrounded by a continuous primary forest and presents a typical Amazonian climate, humid tropical as per the Kopper classification. The human population of the region is comprised of approximately 550 inhabitants, and economic activities include agriculture and livestock.

**Fig 1.**
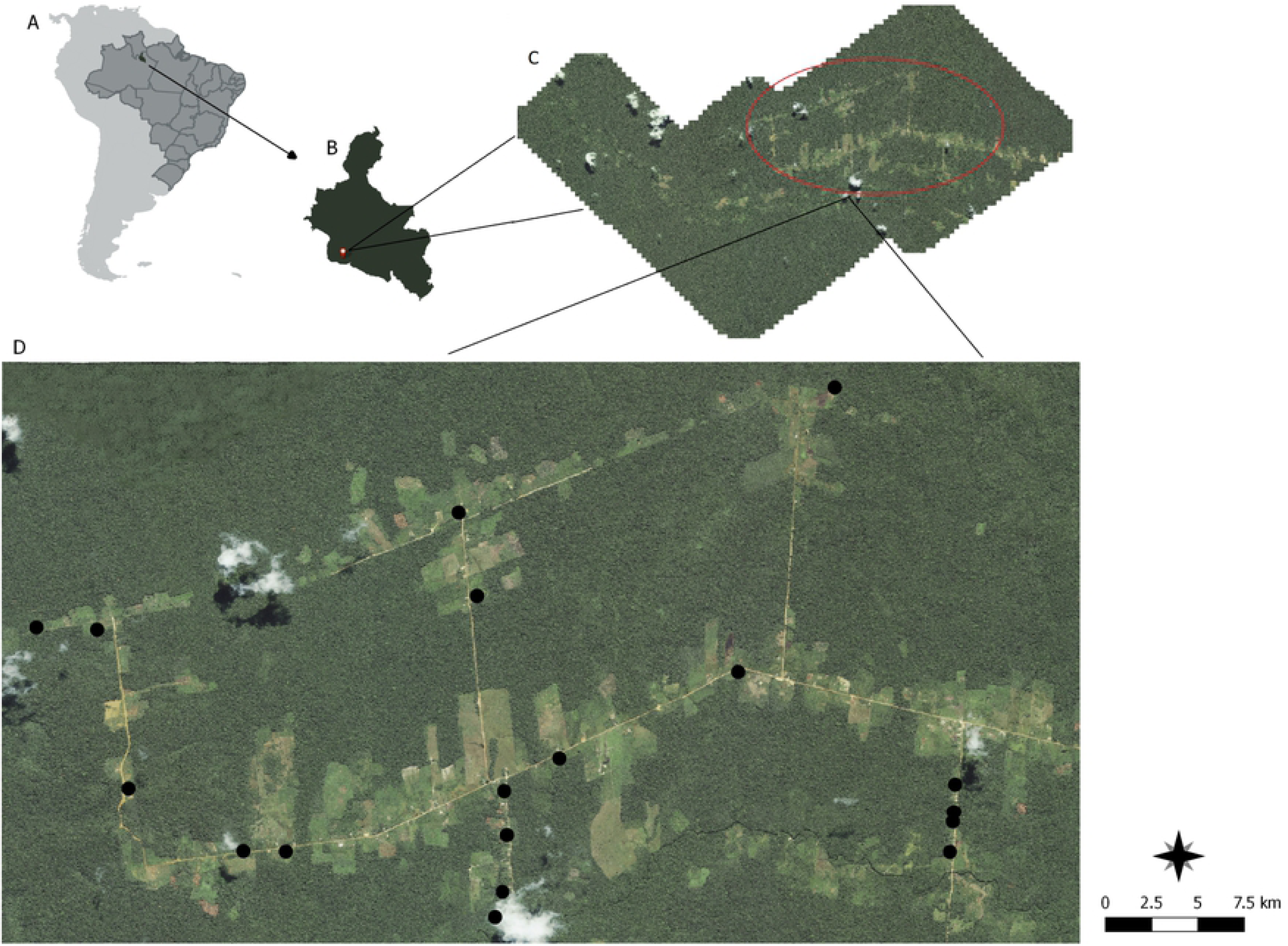
Study area Geographical Information System. A - In different shades of grey; South America, Brazil, and the municipality of Presidente Figueiredo. B - Delimitation area of the Presidente Figueiredo municipality, pointing the agrovillage of Rio Pardo. C - Rio Pardo agrovillage and, in prominence, roads where larvitraps were distributed for immature mosquito collection. D - Location of collection points along agrovillage roads.

The area of the settlement is about 317 km^2^, and includes roads, small villages, wooden houses, gardens, and forest areas. The deforestation rate of the settlement increased from 12.32% in 2008 to 19.79% in 2015.

The delimitation of the settlement area was done through IKONOS™ satellite imagery. Eighteen collection sites were chosen, separated by buffer areas with a radius of 200 m. Sampling points were placed in forest, forest edge, and peridomicile environments.

### Collection of immature mosquitoes

Collections were carried out during four periods of 15 days each, during the months of November 2017 and January and February 2018. For the collection of immature mosquitoes, bamboo internodes, tires, and plastic containers were installed (Fig 2). One unit of each larvitrap type was installed in the forest, forest edge, and peridomicile environments, separated by at least 7 m from each other trap and 1 m from ground level. Nine larvitrap sets were installed at each sampling point area with 162 total traps for each collection event. Traps were filled with 500 ml of tap water.

**Fig 2.**
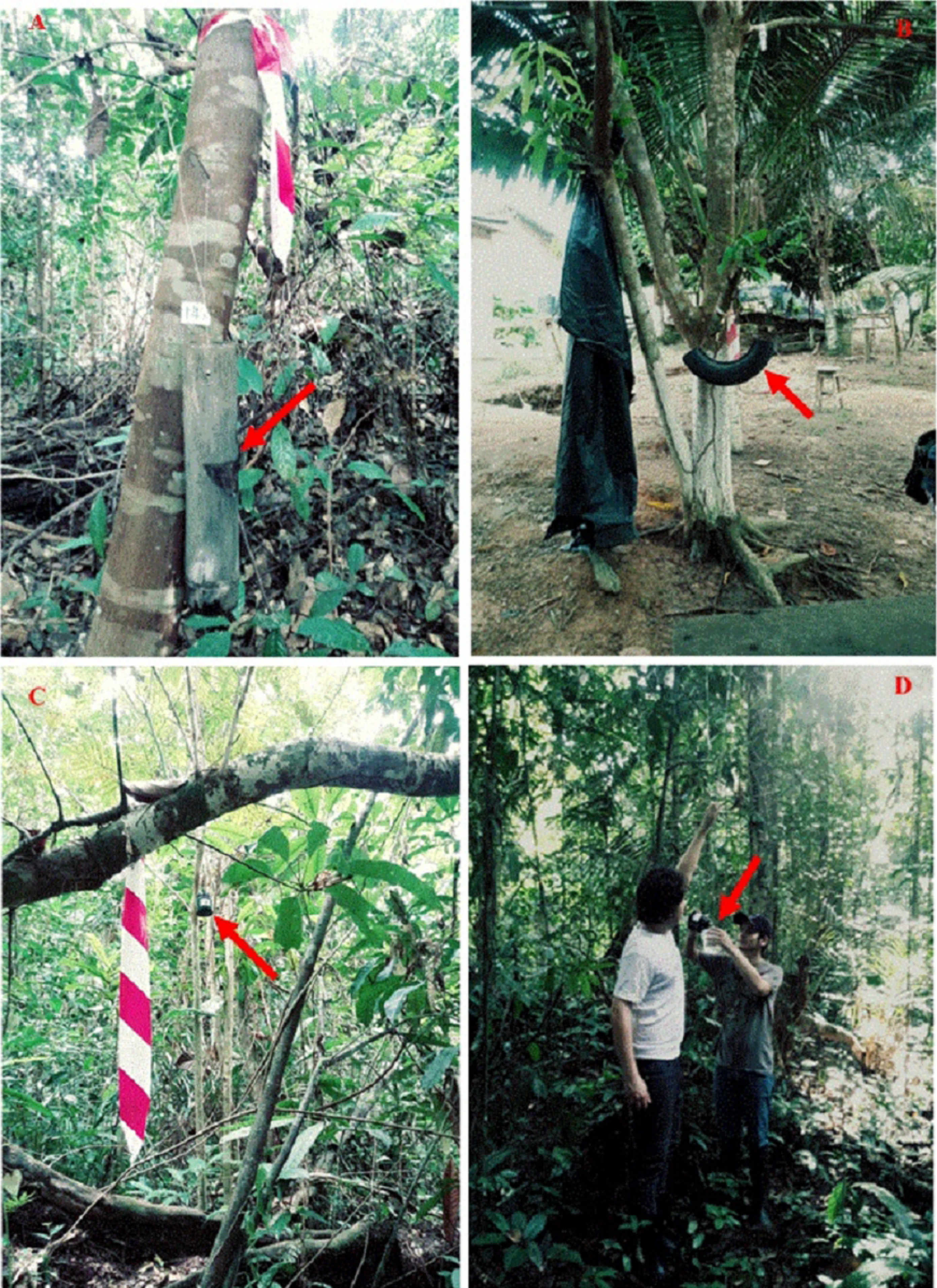
Larvitrap installations in the Rio Pardo agrovillage, Presidente Figueiredo, Amazonas, Brazil. A - bamboo internode. B - tire. C - plastic container. D - material collection. The arrows point to the installed larvitraps for the collection of immature mosquitoes.

The collection of immature mosquitoes began 15 days after installation of the traps. The content of larvitraps was packed in a plastic bag and labelled. Larvitraps were then filled with water again. Collected larvae were sorted to separate them from occasional predators and were reared in plastic containers containing water and organic material present at the time of collection until they reached the fourth instar or adult stage. The collection was made with the permanent license, n° 12186, issued by SISBIO-ICMBio.

Mosquitoes were identified using the identification keys of Lane [24], Forattini [25], Consoli and Lourenço-de-Oliveira [26], and Zavortink [27]. Genera were abbreviated according to Reinert [28].

### Statistical analysis

To evaluate sampling, the rarefaction curve of the identified species was used as a function of the frequency of individuals captured. Shannon-Wiener and Simpson diversity indices, Berger-Parker dominance index, Pielou equitability, and Jaccard similarity were used to analyze the diversity patterns and species distribution between the larvitraps and environments.

To verify the influence of different environments and trap types on mosquito population composition, an Analysis of Permutational Multivariate Variance (PERMANOVA) was performed, which consisted of multivariate non-parametric analysis based on permutations. The Indication Value of Species (IndVal) was applied to verify the specificity and fidelity of the mosquito species to the larvitraps analyzed.

Statistical analyses were carried out using the program Past Version 3.14 [29] and the free statistical software RStudio Version 1.2.1335 [30] with the vegan [31], labdsv [32], ggplot2 [33] and indicspecies [34] packages.

## Results

A total of 10,131 immature mosquitoes were collected, grouped into 10 genera and 20 species. The most abundant species were *Culex urichii* Coquillett, with 2,988 individuals (29.5%), *Trichoprosopom digitatum* Theobald, with 2,746 (27.1%), and *Cx*. (Melanoconion) sp. with 1,052 (10.4%).

The tire was the most effective larvitrap type, with the highest number of individuals, 6,195 (61.1%), followed by the bamboo internode with 2,593 (25.5%) and the plastic container with 1,343 (13.2%). The bamboo internode collected the largest number of species with a total of 17 species, followed by the plastic container with 16 and the tire with 15.

Among the mosquito species collected, *Ochlerotatus argyrothorax* Bonne-Wepster and Bonne was frequently found only in tire larvitraps, *Sabethes cyaneus* Fabricius in the plastic containers, while *Orthopomyia fascipes* Coquillett, *Sa. amazonicus* Gordon and Evans, and *Sa. belisarioi* Neiva were frequently observed in bamboo internode traps. All other species were frequently recorded in at least two larvitrap types (Table 1).

**Table 1.**
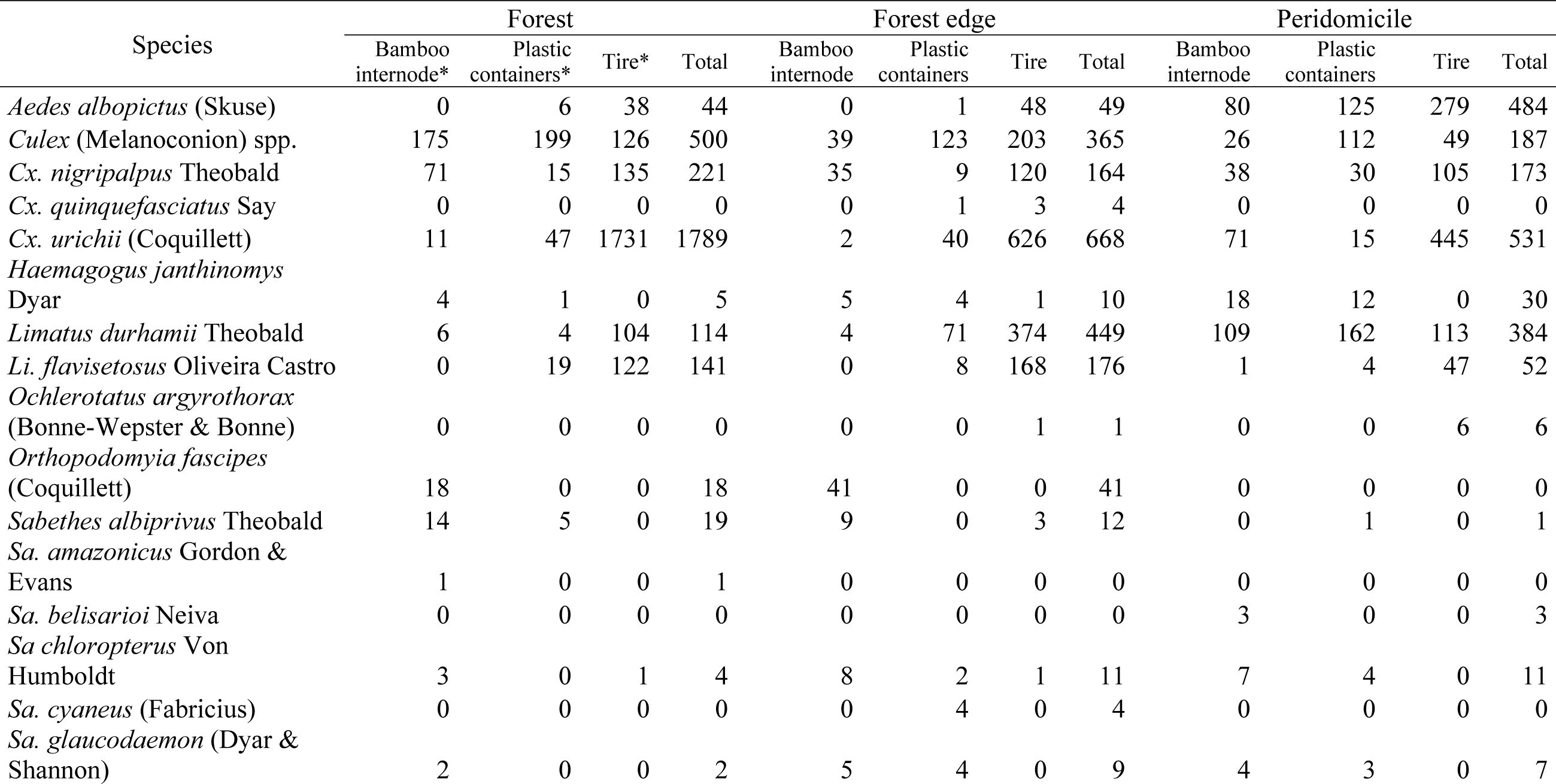
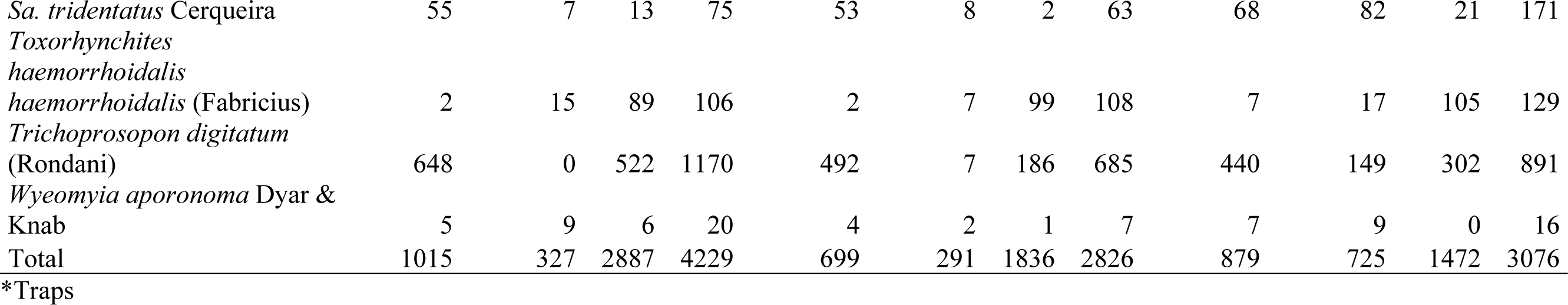
Analysis of captured immature mosquitoes. Distribution of diversity and abundance of mosquitoes found in the bamboo internode, plastic container, and tire traps, located in the forest, at the forest border, or in the peridomicile environment in the agrovillage of Rio Pardo, Presidente Figueiredo, Amazonas, Brazil.

The rarefaction curve of accumulated species richness revealed that plastic container and tire curves had asymptotes, with curves stabilizing around 1,100 individuals and 16 species and 3,500 individuals and 15 species, respectively, in the bamboo internode trap a trend toward stabilization of curves was observed (Fig 3).

**Fig 3.**
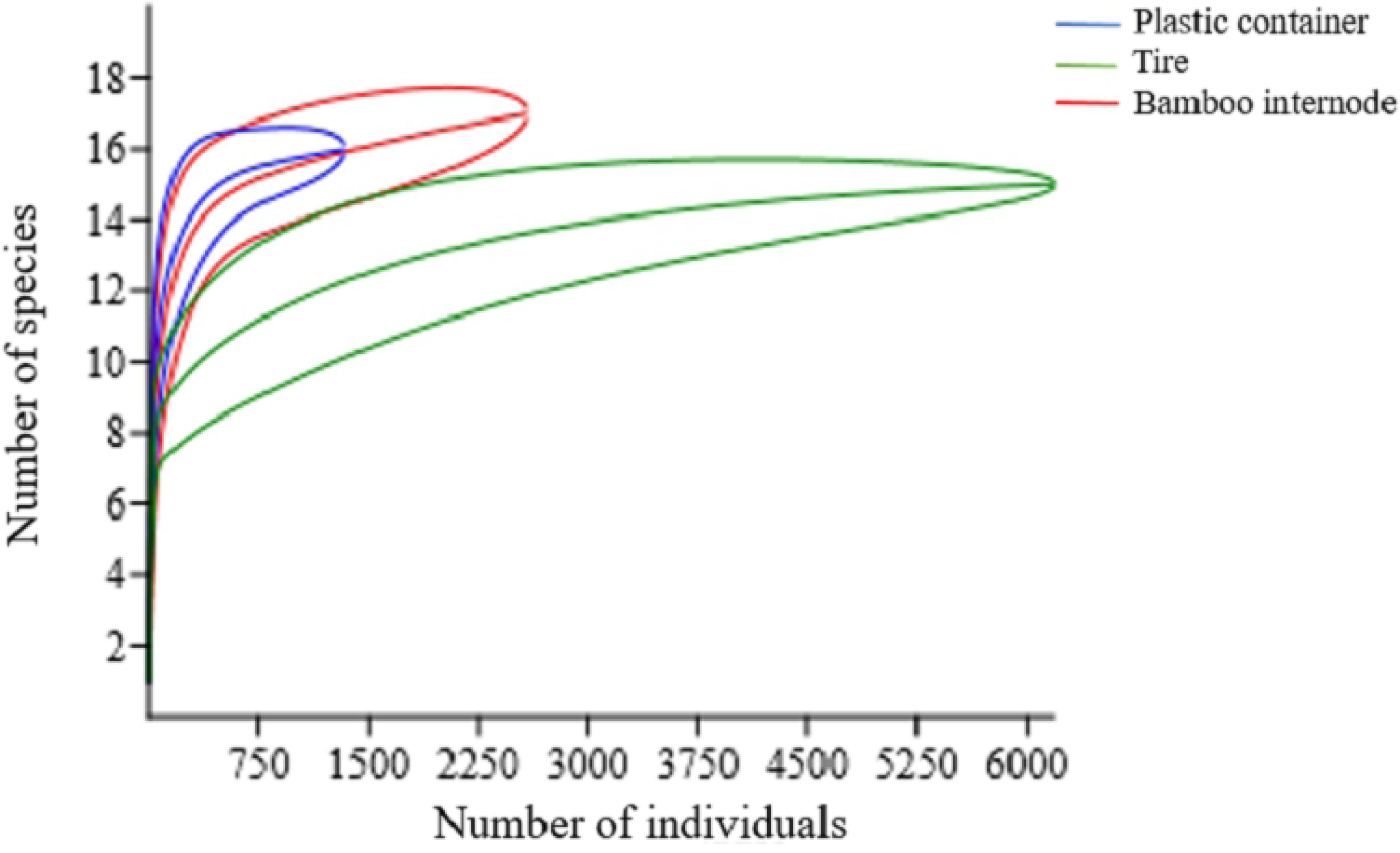
Collected species of mosquitoes. Rarefaction curves representing the accumulated richness of the immature mosquito species collected in plastic container, tire, and bamboo internode traps in the agrovillage of Rio Pardo, Presidente Figueiredo, Amazonas, Brazil periods of November (2017) and January – February (2018).

The plastic container larvitrap presented greater species diversity and equitability of immature mosquitoes when compared to the bamboo internode and tire (Table 2, Fig 3). The greatest similarity in the diversity of immature mosquitoes was observed between the plastic container and the tire (Cj = 0.82).

**Table 2.** Diversity analyses of immature mosquitoes. Ecological indices of species found in larvitraps installed in the agrovillage of Rio Pardo, Presidente Figueiredo, Amazonas, Brazil.

| Ecology attributes | Bamboo internode trap | Plastic container trap | Tire trap |
| --- | --- | --- | --- |
| Shannon - Wiener | 1,527 | 2,07 | 1,741 |
| Simpson | 0,6075 | 0,8268 | 0,7439 |
| Equitability | 0,5389 | 0,7467 | 0,6429 |

The species diversity of mosquitoes was significantly different between the bamboo internode, plastic container, and tire (PERMANOVA pseudo-F=8.84090 p<0.0005). At least 10 bioindicator species were identified according to the IndVal analysis. *Sabethes tridentatus, Tr. digitatum, Sa. albiprivus*, and *Or. fascipes* were the species that presented the greatest specificity and fidelity for bamboo internode larvitraps, while *Tx. haemorrhoidalis haemorrhoidalis, Cx. urichii, Li. flavisetosus, Li. durhamii, Cx. nigripalpus*, and *Ae. albopictus* had the greatest specificity and fidelity for the tire trap. There were no bioindicator species for the plastic container larvitrap (Table 3).

**Table 3.**
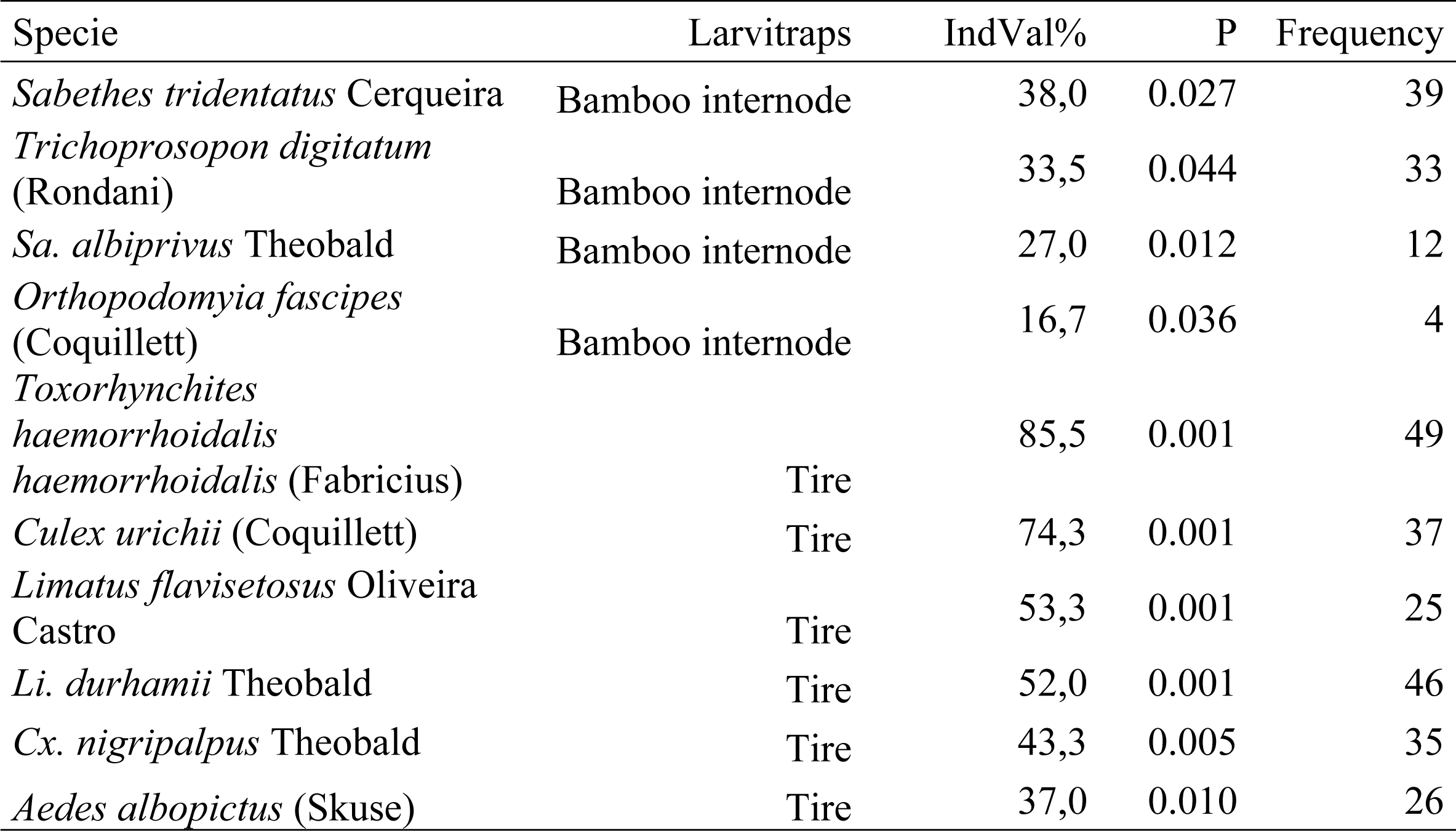
Indication Value of Species (IndVal) of immature mosquitoes. Bioindicator species of bamboo internode and tire larvitraps, according to IndVal, collected in the agrovillage of Rio Pardo, Presidente Figueiredo, Amazonas, Brazil.

Forest, forest edge, and peridomicile environments presented differences in the diversity of immature mosquitoes (PERMANOVA pseudo-F=3.22809 p<0.0010). The forest edge environment had the greatest diversity of species, followed by the peridomicile and forest (Table 4). In all environments, tire larvitraps collected the largest number of individuals, while, in the forest edge, the plastic container and tire larvitraps collected the greatest wealth of species (Table 1).

**Table 4.** Diversity analyses of immature mosquitoes. Ecological indices of species found in larvitraps installed in forest, forest edge, and peridomicile environments in the agrovillage of Rio Pardo, Presidente Figueiredo, Amazonas, Brazil.

| Ecological attributes | Forest |  |  |  | Forest edge |  |  |  | Peridomicile |  |  |  |
| --- | --- | --- | --- | --- | --- | --- | --- | --- | --- | --- | --- | --- |
|  | Bamboo internode* | Plastic container* | Tire* | Environment Forest | Bamboo internode | Plastic container | Tire | Environment Forest edge | Bamboo internode | Plastic container | Tire | Environment Peridomicile |
| Shannon - Wiener | 1,24 | 1,41 | 1,35 | 1,64 | 1,19 | 1,75 | 1,86 | 2,04 | 171 | 2.01 | 188 | 2,03 |
| Simpson | 0,55 | 0,59 | 0,59 | 0,72 | 0,48 | 0,73 | 0,8 | 0,83 | 0.70 | 0.83 | 0.81 | 0,83 |
| Equitability | 0,46 | 0,59 | 0,56 | 0,59 | 0,46 | 0,64 | 0,68 | 0,7 | 0.64 | 0.76 | 0.81 | 0,73 |

207 \* Traps

The diversity of species among larvitraps installed in different environments, presented the following order: forest: plastic container > tire > bamboo internode; forest edge: tire > plastic container > bamboo internode; peridomicile: plastic container > tire > bamboo internode (Table 4, Fig 4).

**Fig 4.**
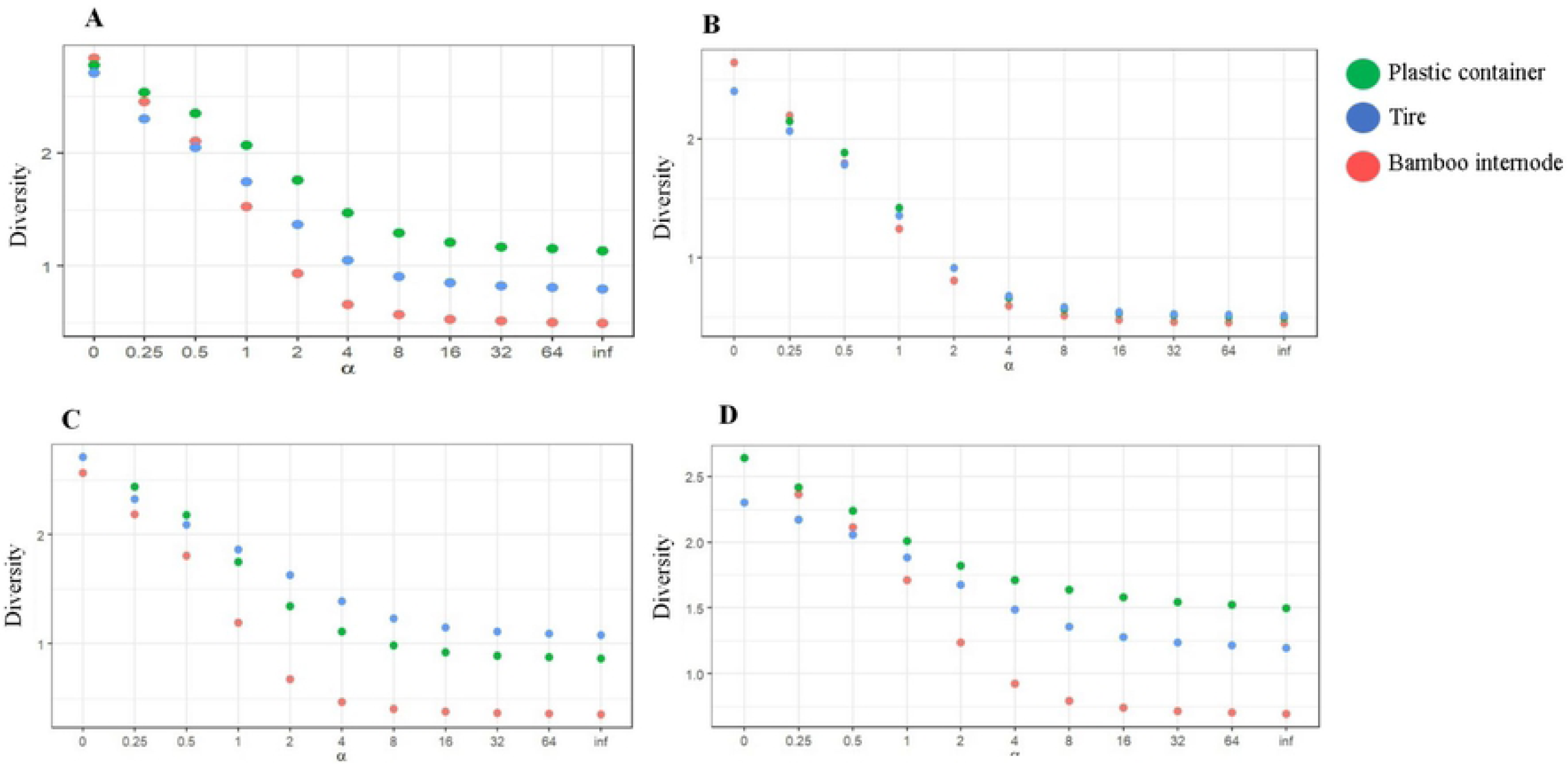
Rényi Diversity profiles of immature mosquitoes. A. Diversity of species found in bamboo internode, plastic container, and tire larvitraps. B. Diversity of species found in bamboo internode, plastic container, and tire larvitraps installed in the forest environment. C. Diversity of species found in bamboo internode, plastic container, and tire larvitraps installed in the forest edge environment. D. Diversity of species found in bamboo internode, plastic container, and tire larvitraps installed in the peridomicile environment. X axis: Alpha (α) zero = log wealth, 1 = Shannon Index, 2 = Simpson Index, Inf = Berger Parker Index. Rio Pardo, Presidente Figueiredo, Amazonas.

The equitability of species in the forest environment was highest in plastic container larvitraps and lowest in bamboo internode larvitraps. At the forest edge, the highest equitability value was observed in tires and the lowest in bamboo internode larvitraps. In the peridomicile, equitability was higher in tire larvitraps and lower in bamboo internode larvitraps (Table 4).

## Discussion

The species of mosquitoes collected in this study represented 7% of the mosquito fauna found in the Amazonas state, which is approximately 270 species [35–44]. Through larvitraps, it was possible to obtain a good estimate of the most common mosquito species present in the Rio Pardo agrovillage, with the exception of the bamboo internode trap. However, it is necessary to use other capture methods, since mosquito species recorded in Rio Pardo so far include 46 species of 17 genera [12].

The high abundance of *Cx. urichii* had not yet been recorded in previous studies of mosquito ecology in the Brazilian Amazon [45,46], however, at low frequencies, this species can be observed in natural and artificial breeding sites in forest areas of the Manaus municipality, Amazonas [47,48]. *Trichoprosopon digitatum*, followed by *Cx*. (Melanoconion) spp., were the second and third species with largest number of specimens. Chaverri et al. [49] also reported the dominance of *Tr. digitatum*, collected in ovitraps in a forest region in Costa Rica. Due to the difficulties of taxonomic characterization of *Cx*. (Melanoconion) species, these were only identified at the section level. However, a high abundance of subgenus species was reported in the studies of Hutchings et al. [39,50] carried out along Amazonian rivers, as well as in the work of Ribeira et al. [51] in areas of the Atlantic Forest of São Paulo.

Larvitrap colonization varied depending on the preference of species. A greater abundance was recorded in tire traps, while the highest diversity was observed in plastic container traps. Similar results of high mosquito abundance in tires were reported by Lopes et al. [13,21], however, these studies also described the artificial tire trap as hosting the greatest diversity of mosquito species when compared to plastic containers, bamboo, and aluminum cans. Lopes et al. [13] also found that the similarity of species was greatest between the tire and bamboo traps, inconsistent with our results, which indicated that the greatest similarity of immature mosquito species was observed between plastic container and tire larvitraps.

The composition of immature mosquito species was different between larvitrap types, as was also observed in investigations of culicid fauna carried out by Calado and Silva [52] and Lopes [21]. Some of the species, as revealed by IndVal, were frequent in breeding sites following patterns commonly described in the literature, as in the case of *Sa. tridentatus, Tr. digitatum, Sa. albiprivus*, and *Or. fascipes*, generally found in natural breeding sites such as the bamboo internode and *Li. durhamii, Cx. nigripalpus*, and *Ae. albopictus*, observed in artificial breeding sites, such as tires [25,27,52–54]. *Toxorhynchites haemorrhoidalis haemorrhoidalis, Cx. urichii*, and *Li. flavisetosus* have not yet been associated with tires as their breeding site.

The greatest diversity of immature mosquitoes species found at the edge of the forest suggested a change in the habitat of these populations, and similar results were reported in the studies of Steiger et al. [55]. Chaverri et al. [49], in their study of immature mosquito fauna in different environments, observed no difference in mosquito populations between the primary and secondary forest environment. Ribeiro et al. [51] suggested that environmental stresses increase the number of niches favorable to mosquitoes and, thus, promote the greater diversity of mosquitoes in anthropogenic environments.

Tire larvitraps, when compared to other types present in the forest, forest edge, and peridomicile environments, had the highest species abundance in all environments, as well as the greatest wealth of species in the forest edge environment. It is believed that species that develop in tires originally breed in holes of trees and have a preference for tires due to their similar characteristics, such as a dark environment, shading, presence of organic matter retained inside, retention of excessive water volume, and a slow evaporation process [21,56,57]. According to Beier et al. [57], some species ended up dominating the tire trap, and, in the case of this study, the dominant species was *Cx. urichii*. Rubio et al. [58], evaluated the success of tire colonization across a gradient of less and more urbanized areas in Argentina, and they found that the majority of species captured frequented both areas. In addition, there was a trend of increasing abundance of vector species in less urbanized environments.

In the forest edge environment, culicid species found in plastic container larvitraps had a species richness similar to that of mosquito populations in tire traps. However, the fact that there was a low abundance of 10 species present in the forest edge tire larvitraps, with values between one and nine individuals per species, may suggest that these species preferred the plastic container in this environment for opportunistic colonization, since the forest edge did not provide a large variety of natural breeding sites as the forest would.

In general, in all the studied environments, artificial larvitraps, namely the plastic containers and tires, had higher immature mosquito species diversity and equitability values when compared to bamboo internode larvitraps.

Although there is a great diversity of breeding sites and different patterns of choice for oviposition site, which vary between mosquito species, artificial larvitraps presented the best conditions for the development of immature mosquitoes, independently of the environment. Thus, it is plausible to suggest that some of the species collected in the current study demonstrated a certain eclecticism at the time of choosing the place of oviposition and had success in the development of their larvae in artificial breeding sites. In addition, the smaller diversity of species found in the bamboo internode larvitraps in all environments reinforces the idea that only a few species, especially those considered wilder, such as the Sabethini tribe, have a preference for small natural breeding sites found in wild environments [25,59].

All vector species found in this study were collected in plastic container and tire larvitraps, with a greater abundance in the forest edge and peridomicile environments. These observations suggest that such artificial breeding sites may be risk factors for infection with typical wild arboviruses [53,60].

Lopes [20] actively searched for immature mosquitoes in artificial breeding sites in rural areas of Londrina, Paraná, and showed that the abundance of immature mosquitoes was higher in boxes d’agua, followed by tires and troughs. In addition, species *Cx. quinquefasciatus* and *Li. durhamii* were found in all artificial breeding sites, and, thus, Lopes and colleagues suggested that these were species adapted to anthropogenic environments.

Through this study it was possible to identify wild Amazonian species that colonize larvitraps located in environments with different degrees of anthropization. The preference for artificial larvitraps observed in the current study may be associated with the opportunism of females or may indicate a change in the habits of these wild species, including vector species. However, further studies are required to assess these behaviors.

## Acknowledgments

We thank Fernanda Fonseca MSc for providing data on the territorial delimitation of the Rio Pardo settlement, Dr. Bernardo Horta and Antônio Balieiro MSc for their assistance in the statistical analysis, Eric Marialva MSc, Antônio Leão MSc, Ricardo Mota, and Sebastião Dias for helping with field collections, and Gervilane Ribeiro for mosquito class identification. In memory of Patrícia Dantas, we would like to thank her for helping to make the traps and growing the immature mosquitoes in the laboratory. We thank for the grant of the scientific initiation scholarship of the co-author of the article, Heliana Belchior n° 00812016 - PAIC - AM – 2016. We would also like to thank Fiocruz / VPEIC for the scholarship used to carry out the study and the post-graduate course in Condições de Vida e Situações de Saúde na Amazônia of the Instituto Lebônidas e Maria Deane - Fiocruz Amazônia, for the training and qualification of Master in Public Health. We would like to thank Editage (www.editage.com) for English language editing.

## Supporting information

S1 Table. **Data from mosquito collections, carried out in Rio Pardo, Presidente Figueiredo, Amazonas, Brazil**.

